# MicrogliaST: a web server for microglia spatiotemportal pattern analysis in normal and disordered brains

**DOI:** 10.1101/2022.01.08.475469

**Authors:** Xiaoling Zhong, Feng Li, Guiyuan Tan, Li Yi, Jiaxin Zhao, Wanqi Mi, Yu Zhang, Congxue Hu, Xia Li, Yingqi Xu, Chunlong Zhang

**Affiliations:** College of Bioinformatics Science and Technology, Harbin Medical University, Harbin 150081, China; Center of Cerebrovascular Disease, Zhuhai People’s Hospital, Zhuhai hospital affiliated with Jinan University, Zhuhai 519000, China

**Keywords:** microglia, spatiotemporal analysis, single cell RNA seq, glioma, Alzheimer’s disease

## Abstract

Brain is the most complex organ of living organisms, as the celebrated cells in the brain, microglia play an indispensable role in the brain’s immune microenvironment. Microglia have critical roles not only in neural development and homeostasis, but also in neurodegenerative diseases and malignant of the central nervous system. However, little is known about the dynamic characteristics of microglia during development or disease conditions. Recently, the single-cell RNA sequencing technologies have become possible to characterize the heterogeneity of immune system in brain. But it posed computational challenges on integrating and utilizing the massive published datasets to dissect the spatiotemporal characterization of microglia. Here, we present microgliaST (bio-bigdata.hrbmu.edu.cn/MST), a database consisting of single-cell microglia transcriptomes across multiple brain regions and developmental periods. Based on high-quality microglia markers collected from published papers, we annotated and constructed human and mouse transcriptomic profiles of 273,374 microglias, comprising 12 regions, 12 periods and 3 conditions (normal, disease, treatment). In addition, MicrogliaST provides multiple analytical tools to elucidate the landscape of microglia under disorder conditions, conduct personalized difference analysis and spatiotemporal dynamic analysis. More importantly, microgliaST paves an ingenious way to the study of brain environment, and also provides insights into clinical therapy assessments.

## Introduction

Parenchymal microglia was major component of myeloid cells involved in heterogeneous Central Neural System (CNS) immune micro-environment [1-3]. Microglia are multifunctional cells that interact with numerous other cells in the CNS, including neurons, astrocytes, and oligodendrocytes, showing its key biological roles in brain development, homeostasis, and disorders. Recently, many lines of evidence indicate that dysregulation of microglia functions contributes to the pathogenesis of neurodegenerative diseases, such as Alzheimer’ s disease [4-6], as well as neurodevelopmental and psychiatric disorders [7, 8]. In addition, microglia play important roles in the injured brain [9] and brain-related tumors, such as glioma [10]. In the meanwhile, spatially and temporally restricted microglia characterization in the central nervous system during development or disease was necessary. And a single-cell analysis of CNS tissues during homeostasis in mice revealed time- and region-specific subtypes of microglia [11]. Furthermore, specific brain regions or developmental stages were found to be involved in different diseases [12]. Although the evidence is accumulating, such studies are still very limited on the number of microglia and kinds of disease types. To better understand the role of microglia in brain disorders, there is a pressing need for a comprehensive exploration of microglia’s spatiotemporal characterization in brain tissue.

With the wide application of single cell RNA sequencing (scRNA-seq) in transcriptomic profiling, plenty of scRNA-seq related databases have been developed. Among them are comprehensive databases and other disease-specific databases. In terms of comprehensive databases, there exist Single Cell Portal, PanglaoDB [13], Single Cell Expression Atlas [14], Human Cell Atlas Data Portal [15], SCPortalen [16], SCDevDB [17], JingleBells [18], STAB [19] and so on. Most of these databases classify a mass of single-cell expression datasets from both healthy and diseased tissues involving human and mice. In general, these databases contain only basic analyses of cell clusters and differential gene expression profiles, and do not contain in-depth analyses of disease types or clinical information. Take STAB for example, although it focuses on clinical information, especially the spatiotemporal characteristics of the brain, it does not address disordered brain. Although these databases are relatively targeted, some of these do not include microglia, such as SC2Disease [20], CancerSCEM [21] and TISCH [22]. Other database, like CancerSEA, only focuses on the correlation between genes and these cancers, but does not provide expression information for each gene in specific cell types and tissues[23]. In summary, the integration of these cross-platform single-cell datasets, accurate cell type identification, and comprehensive online analysis platforms are still insufficient, leaving great challenges for microglia research.

Given the important role of microglia in the regulation of brain homeostasis, immune response, and development, there is an urgent need for a comprehensive multi-species microglia atlas, which is important to better understand the differences between microglia molecular characteristics, dynamic changes, and the difference between disease and normal. Here, we present microgliaST, a comprehensive database for microglia spatiotemporal characterization in normal and disordered brains. By collecting single-cell transcriptome data from the publicly available sRNA-seq dataset and analyzing it using standard procedures, MicrogliaST classified microglia from 12 brain regions and 12 developmental stages across both human and mouse species. This database presents a comprehensive cellular landscape of microglia and provides user-defined differential expression results for researchers of brain disease.

## Materials and Methods

### Data collection and organization

We collected publicly available human scRNA-seq data as well as their metadata by searching PubMed and the public databases, Gene Expression Omnibus (GEO) [24]. The single-cell or single-nucleus RNA sequencing datasets were retrieved with the following keywords: (i) single-cell RNA seq, single-cell transcriptome and single-nucleus seq, in combination with (ii) brain, central nervous system, brain disorder, glioma, Alzheimer’s disease and the name of specific brain regions. Each dataset was then manually confirmed and curated. The datasets were further required to be generated from brain tissues from human and mouse species without considering cell lines or brain organoids. As for the following integration, only the raw count format was considered. To investigate the dynamics of brain cellular compositions and gene expression from embryonic to aging, we further assigned samples to distinct developmental periods as described by Kang et al.[25, 26]. As a result, 12 brain regions and 12 developmental periods for human and 12 brain regions and 12 developmental periods for mouse were considered. The detailed data selection workflow was provided as **Figure 1**.

**Figure 1.**
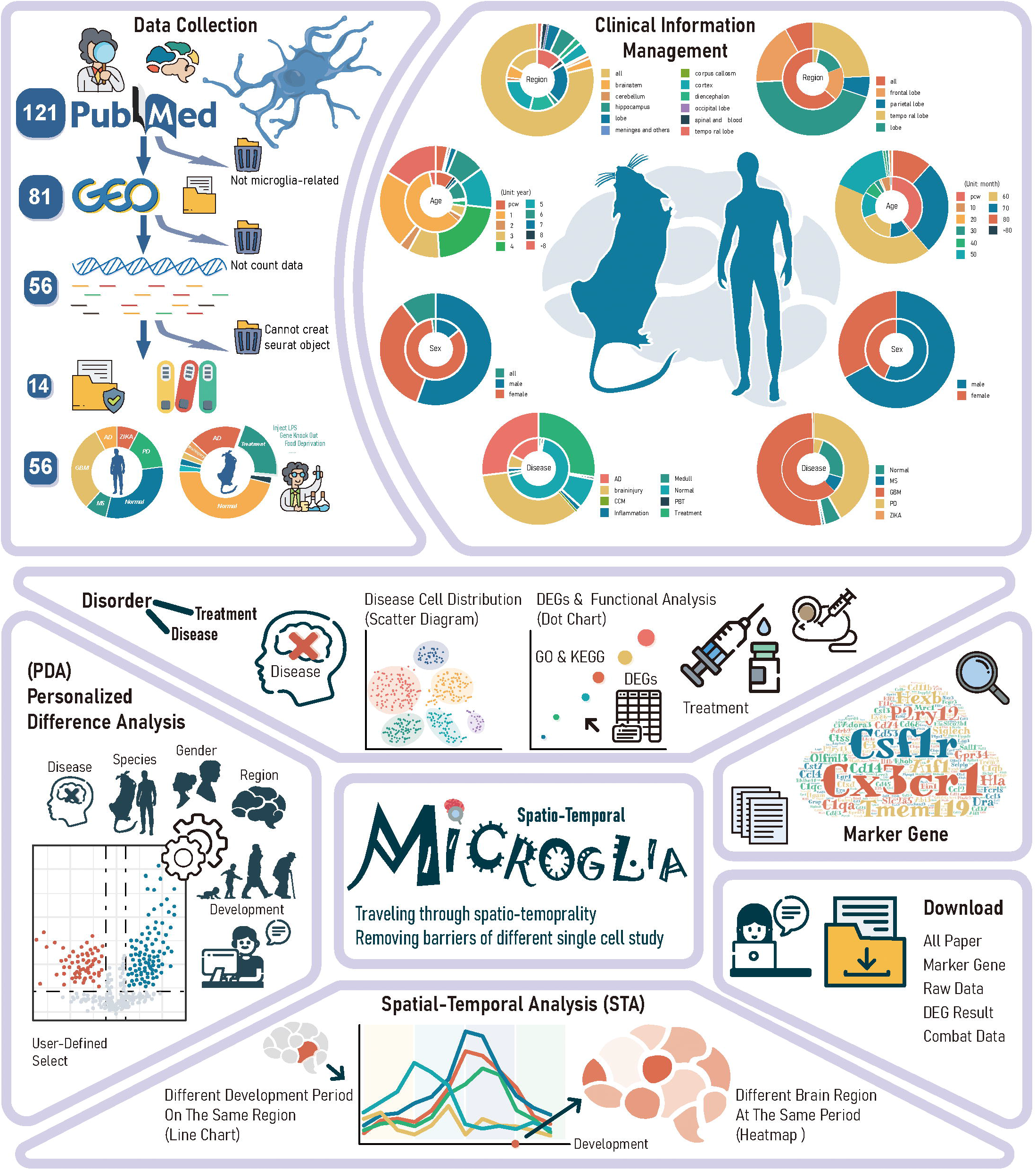
The framework for screening microglia related scRNA seq dataset.

### Quality control of scRNA-seq dataset

For each dataset, we firstly applied the R package Seurat [27] to transform the raw count data into Seurat objects, and then removed mitochondrial and ribosomal related genes according to the standard procedure. In order to remove the empty droplets, low-quality cells and other interference factors that may interfere with our results, we furthermore removed those unqualified cells according to the following indicators: i) The number of genes expressed in each cell must >200, ii) the total count of each gene must >1000. For each gene in a cell, its expression level was normalized and defined as the ratio between its counts and the total counts of genes expressed in the cell, which was then multiplied by a scale factor (10,000 by default) and log-transformed.

### Annotation of microglia

To in silico select microglia cells from scRNA-seq dataset, we reviewed and collected the microglia cell markers in 66 associated literatures, and regarded the consistent markers which existed at least three literatures as the high-quality microglia markers (see **Table S1**). For each scRNA-seq dataset obtained above, which undergo filtering and quality control of standard process, the original mixed brain cells were divided into clusters by the adjusted uniform parameters for dimensionality reduction and clustering. Subsequently, we used the function findAllmarker in the Seurat package to identify the top 100 differential genes in each cluster, and conducted a hypergeometric test between these differential genes and the high-quality microglia markers. Clusters with P-values less than 0.05 were selected as the result of manual annotation of microglia clusters from each datasets.

### Integration of datasets and Batch effect removal

After quality control and annotation, data from the same species and the same condition (normal or certain disorder) are integrated into a merged matrix by the merge function in the Seurat package, and its batch effect was further removed by the Combat function in the SVA package. After that, we integrated data sets from different studies systematacially, making cross-dataset analysis possible, which is conducive to the full utilization of brain scRNA-seq datasets.

### Data availability and downloads

All pre-processed data and curated metadata are available for download in comma-separated flat file format. Visualizations can be downloaded as PNG or HTML files that preserve the same interactivity as available from the web application, tables can be downloaded in comma-separated value format.

### Database construction and implementation

MicogliaST was implemented in Java with SpringBoot, Ngnix, Tomcat, Mybatis and MySQL for back-end data interaction, and React for the front-end display. Both visualization and statistical analyses were performed with R, where the gene expression profiles were visualized with the R-package ‘plotly’. The association of a query gene list with a certain brain condition (normal or disorder) was performed with enrichment analysis of the genes over the query gene set, where one-tailed Fisher’s exact test was adopted here with genes expressed in >10% cells as background. With a userfriendly interactive interface, the users can easily browse and query their gene expression profiling across brain regions and developmental periods.

## Results

### Overview of microgliaST

microgliaST is a comprehensive single-cell database specialized in microglia-related diseases, covering multiple brain regions and development stage and showing the dynamic changes of gene expression during the lifespan through a series of visualization methods. For mouse, 11 regions, 10 development period, 8 condition (including multiple treatments) were obtained. And for human, 4 regions, 10 development period, 5 condition (including GBM and PD) were obtained (see **Figure 1**).

The “Disorder” page shows the comparison between microglia-related disordered samples and normal samples, which is divided into two main panel: Disease and Treatment. The “Disease” and “Treatment” pages present the information about certain disorder type you choose at the beginning. Then, there are the distribution of microglia cells according to clinical information, a difference expression gene list between normal samples and this disease. Moreover, base on the list of selected genes, the enrichment analysis of these genes can also be generated. A variety of microglia-related brain diseases are included in our database, containing both human and mouse, which is of great practical significance for clinical research

The ‘PDA’ page is a particular tool to conduct differential expression analysis for researchers in various fields. “PDA” means Personal Differential Analysis, give users a very personalized option in terms of calculating differential genes. Users can freely choose two expression matrices according to five terms such as gender, brain region, time and so on. And further explore the differences in gene expression that they were interested in

The “Dynamic Analysis” pages show the trend variation of gene expression of different brain region or development periods. Given a gene list (less than 5) and other constraints, the expression dynamics of these genes for each regions/periods will be displayed.

### Data quality organization

In order to evaluate the quality of our annotated microglia, we compared the results obtained from manual annotation with the microglia results presented in the original literature. Compared with the cell type labels described in the original studies from which the datasets were retrieved, the cell types defined by our cellular clusters had a concordance rate of 92.3% (see **Figure S1**), indicating the confidence of our cell types defined here.

### Database function

#### 1. Personalized Differential Analysis

PDA (Personalized Differential Analysis), is a highly alternative and flexible tool conducting differential expression analysis for researchers in various fields. Users can performed personal difference expression analysis according to 5 terms below: species, gender, brain region, time and sex. The steps of how to select the matrix used to make difference are as follows: (1) Choose which species (human or mouse) you want to explore. (2) Choose the only options you want to conduct difference analyze (Gender, Region, Time, Disease). (3) Select any secondary options between remaining three options that you want to explore (multiple options are available). (4) Click the submit to start the difference analysis. If you have no idea how to choose, you can click example to see the official recommendations. PDA is the most comprehensive tool to analysis microglia’s differential expression at the spatiotemporal level, enabling users to compare gene expression across species and data sets. All data sets are processed, annotated, and manually managed to remove the barriers of different data sets and different studies, and to connect spatiotemporal areas that are not together, which is conducive to cross-study comparison and data reuse.

#### 2. Marker Gene

On home page, the users can search the marker gene we want to explore directly, or we can just click on the word cloud to obtain information about corresponding gene. And, we offer a Marker Gene panel specially. In this panel, we provide all the relevant information about marker genes collected by manual retrieval. The charts and images on this page show an overall overview of the frequency and expression of all marker genes in all datasets.

#### 3. Disorder Exploration

On this page, user can obtain basic information about the selected disease, and explore the disease according to different species. Besides, differential analysis and gene-set enrichment analysis can also be performed. Take AD for example, the first thing that comes into sight is a basic introduction of AD disease extracted from Wikipedia. Next we can select human or mouse to see the distribution of cells in our database, and we provided several ways to show the single-cell sequencing datasets by reducing dimensions using PCA, UMAP and TSNE methods. And then is the list of differential expressed genes for corresponding dataset, and user can select the list of differential expressed genes by setting different cut-offs, 0.05 or 0.01. In the meantime, GSEA results corresponding to these differential genes will be generated below the gene list. Users can explore the functional enrichment results of these differential expressed genes on our page, and we also provide a download channel for the list of differential expression genes in a personalized and flexible way.

#### 4. Spatio-Temporal Analysis

On this page, users can explore the dynamic expression pattern of interesting genes under spatial and temporal dimension. Firstly, we choose the items we want to explore (Time or Region). Next, we choose the disease, species and development period/brain region we wish to research. Last, we can enter the genes we want to explore (less than 5) and click on submit to check and compare the expression patterns of these genes. Under the spatial and temporal distribution of cells, we can explore the dynamic variation trend of gene expression in single cell and study the microglia cells from a developmental perspective.

#### 5.Data Acquisition and Submission

In order to facilitate the use of related studies, we provide a large number of related files to download, including raw data download from GEO, expression matrix for all microglia cells extraction through our process, all dataset with batch effects removed and all references. In addition to download, we also provide the data submit function. Users can submit latest microglia single cell sequencing transcriptome datasets in the submit page. We will check the entries and add it in our database, as well as contact the submitter as soon as possible.

## Discussion

MicrogliaST is a comprehensive resource of microglia in brain, covering almost all brain regions and developmental periods in both human and mouse species. Through standardized and consistent classification criteria, we reused the clinical information of microglia, so that the spatiotemporal characteristics can be clearly presented. Furthermore, as microglia are important immune cells in the brain, playing an important role in development, homeostasis, immune response and other aspects. We also provide users with a flexible differential analysis tool and a concise disease profile page, so that clinical researchers without a background in bioinformatics can easily explore microglia-related explorations. In conclusion, our database provides multiple channels to explore microglia, and abundant downloadable resources expand a new horizon for researchers in all professions, promote the collaboration between clinical and bioinformatics, make cross-dataset analysis and clinical application possible.

We also noticed that our database still had some flaws: (1) The variety of diseases is not rich enough; (2) Since clinical information is hard to acquire, we don’t have enough information to use. Given all this, we will replenish more data sets in other formats (including Bulk data, chip data, etc.), dig into the clinical information behind these data sets, and strive to enrich our database in the following work.

## Declarations

## Ethics approval and consent to participate

Not applicable.

## Consent for publication

Not applicable.

## Acknowledgements

Not applicable.

---

**Figure S1.**
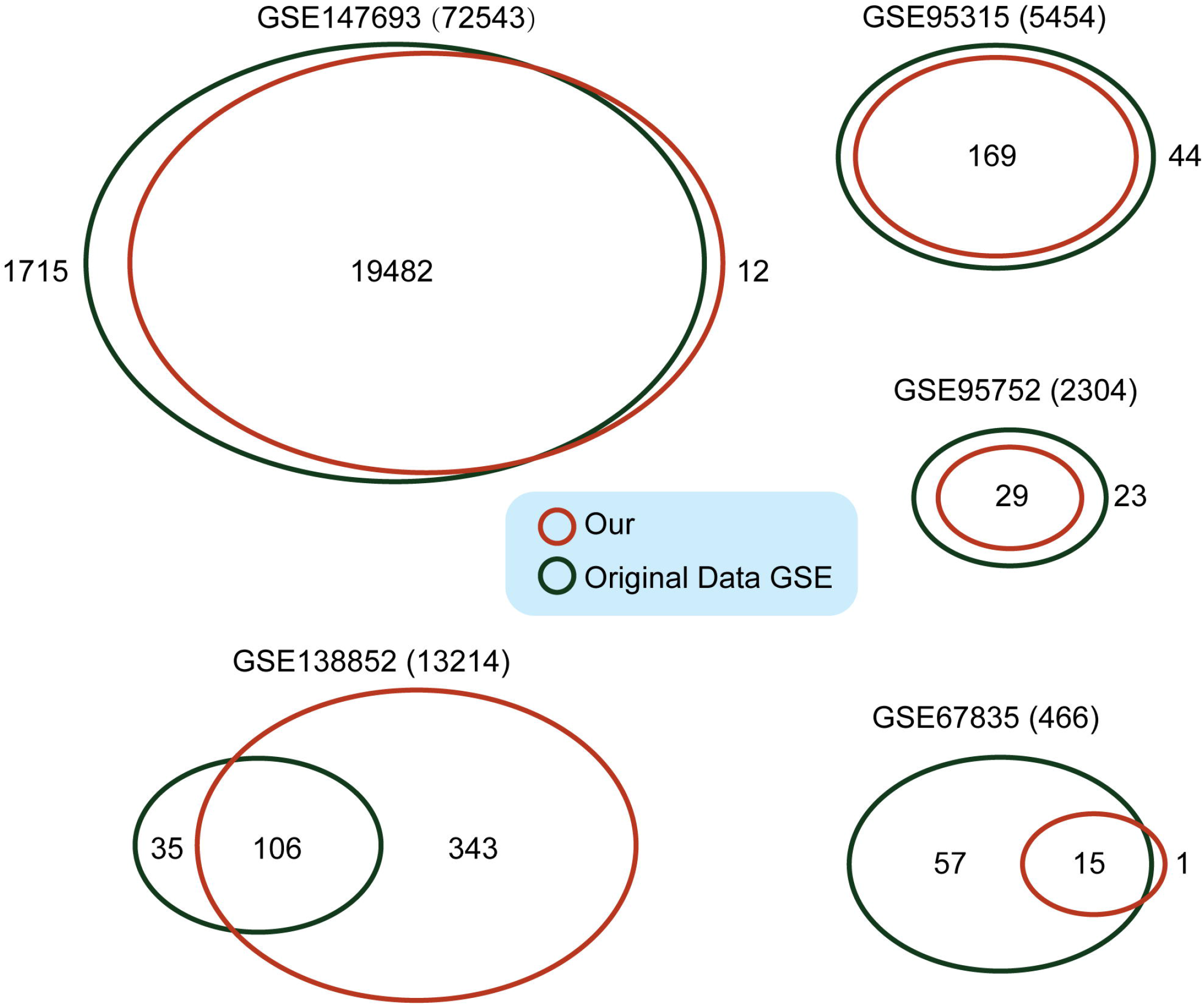
The comparition between the results obtained from manual annotation and the microglia results presented in the original literature

